# Systematic characterization of the yeast secretome under diverse proteosynthetic stress conditions reveals secretion of functional ER chaperone BiP

**DOI:** 10.64898/2026.05.21.727034

**Authors:** Shulei Liu, Benjamin L. Schulz

**Affiliations:** School of Chemistry and Molecular Biosciences, The University of Queensland, St Lucia, 4072, Australia

## Abstract

The yeast secreted proteome plays critical biological roles and influences product and production parameters in industrial fermentation. Systematic profiling of the response of the yeast secretome to intrinsic and extrinsic factors is therefore essential for understanding these functions and for optimizing manufacturing processes. Here, we characterized the yeast secretome under diverse proteosynthetic stress conditions, including glycosylation deficiency, oxidative, reductive, and thermal stresses. The secretome was predominantly composed of conventionally secreted proteins, while a subset of proteins appeared to be secreted via unconventional pathways. Distinct secretome profiles were observed in response to different stressors, driven by a combination of altered intracellular proteomes, altered canonical secretion, and altered cell lysis and unconventional protein secretion, while reflecting the underlying metabolic state of the cells. Heat stress did not impact protein glycosylation but did cause similar protein misfolding stress to N-glycosylation deficiency. Intriguingly, canonically intracellular chaperone BiP was abundant in the secretome in particular stress conditions where its activity would be beneficial. BiP interacted with probable extracellular client proteins *in vitro*, consistent with it acting as a functional extracellular chaperone/holdase in conditions such as reductive stress in which client proteins could be misfolded outside the cell.

## Introduction

*Saccharomyces cerevisiae* is the most widely-used fungal model organism and is a versatile cell factory for biopharmaceutical, fuel, food and beverage production^1,2^. In all of these applications, wanted and potentially unwanted host cell proteins (HCPs) will also be produced and secreted by the yeast cells into the growth media, impacting the purity, quality, and efficacy of the product^3^. Removal of cell-derived contaminants such as HCPs is a major challenge of downstream processing during precision fermentation or biopharmaceutical production, and accounts for a large part of production costs^4^. Yeast proteins are also secreted into the fermentation mixture during beer and wine production, and remain in the final product, affecting their quality and properties^5,6^.

The secretome, the proteins actively secreted or released by yeast cells into their surrounding environment, plays crucial biological roles for yeast^7^. These secreted proteins include digestive enzymes such as proteinases, cell wall remodelling enzymes such as glucan synthases, and stress-responsive proteins such as seripauperins^8-10^. Conventional protein secretion in eukaryotes starts in the endoplasmic reticulum (ER) driven by recognition of an N-terminal signal peptide that triggers translocation of the nascent polypeptide into the ER lumen, protein folding in the ER, trafficking through the Golgi, and ending with sorting and secretion of mature proteins at the plasma membrane^11^. Alternatively, secretory proteins lacking signal peptides can still be exported through unconventional protein secretion (UPS) pathways^12^. Such proteins are also known as ‘moonlighting’ multifunctional proteins, and include glycolytic enzymes, chaperones, and translation factors^13^. UPS also can be triggered by ER dysfunction or by stress that perturbs the canonical ER/Golgi-dependent secretion pathway. For some proteins secreted via UPS pathways under normal condition, cellular stresses may enhance their release^14^. Numerous non-canonically secreted proteins have been detected in the secretome of both yeast and mammalian cells, including Enolase, Galectins, Hsp70, and Pgk^15-21^. The high similarity in unconventional secreted proteins across species suggests that UPS pathways are evolutionarily conserved and physiologically important. In the canonical secretion pathway, the capacity of the ER to fold and process secretory proteins is not unlimited. ER stress occurs when the capacity of the ER to fold proteins is overwhelmed, leading to the accumulation of unfolded proteins in the ER lumen^22^. This accumulation can be triggered by genetic or environmental perturbations altering ER homeostasis, such as mutations in protein-coding genes and oxidative stress^23,24^. To adapt to these stresses, the unfolded protein response (UPR) is activated as an intracellular signalling pathway to transmit signals of the protein folding status of the ER lumen to the cytoplasm and the nucleus, consequently triggering a series of adaptive mechanisms including upregulation of ER chaperones, expansion of the ER, and activation of ER-associated degradation (ERAD) to alleviate the stress states^25,26^. UPS pathways can also be subject to regulation, with cellular stress conditions including heat shock, osmotic stress, starvation, inflammation, and altered autophagy being reported to induce or stimulate unconventional secretion of certain proteins in yeast and/or mammals^14,15,21,27-31^.

Proteins secreted from yeast cells are present in the products of industrial fermentation processes, and so any changes in the process or in yeast physiology may change the yeast secretome and impact product quality or downstream processing^32^. Therefore, understanding the global yeast secretome landscape under different cellular stress conditions relevant to yeast physiology or industrial bioreactor parameters can provide valuable insights into the fundamental biology of yeast cells and their interactions with their environment, with opportunities to improve the efficiency of industrial fermentation bioprocesses. Here, we monitored the yeast secretome in response to diverse intrinsic and extrinsic stresses and linked these responses to physiological adaptation to the specific applied stress. We identified the ER resident chaperone BiP in the secretome of cells in a subset of stress conditions, and investigated its role as an extracellular chaperone/holdase to stabilise misfolded extracellular proteins in these stress conditions.

## Results and Discussion

### Growth pressure induced by proteosynthetic stressors on yeast

Proteosynthetic stress can be triggered by external or internal perturbations to the cell that interfere with productive protein folding in a variety of ways. These stressors can be particularly problematic for secretory proteins which often undergo complex post-translational modification processes, including glycosylation and disulfide bond formation/isomerisation in the ER. To understand the secretory response to various stresses, we selected diverse types of stressors and titrated their dose to yeast cells: organic solvent (ethanol; EtOH), reductive (dithiothreitol; DTT), heat, *N*-glycosylation (tunicamycin and Δ*ost3*), and *O*-glycosylation (Δ*pmt1*) (Table S1). To compare the responses of yeast to these diverse stress conditions, we first assessed the impact on growth of different levels of each stress. We aimed to choose stress conditions that resulted in equivalent and appropriate perturbations from each stressor that slightly perturbed yeast growth. We found that equivalent and slight growth perturbation relative to unstressed wild type (WT) yeast could be achieved with 20% ethanol, 3 mM DTT, 0.3 μg/mL tunicamycin, and 37 ℃, as well as the gene deletion strains Δ*ost3* and Δ*pmt1* (Fig. S1, Table S2).

### Yeast secretome profiling of the response to different stress conditions

To comprehensively profile changes to the yeast secretome in response to the diverse stress conditions we used quantitative data-independent acquisition (DIA) liquid chromatography with tandem mass spectrometry (LC-MS/MS) proteomics. As expected, the overall secreted proteome was modest in complexity, with 165 proteins identified across all conditions (Supplementary Data 1-4), with many proteins only identified in a single condition (Fig. 1A). No proteins were uniquely identified from unstressed WT yeast; proteins that were only measured in a single condition were instead identified from specific stress conditions. Most proteins identified in the secretome of WT yeast were conventionally secreted proteins from the canonical secretory pathway such as cell wall proteins Ecm33 and Pst1, and cell wall remodelling enzymes Gas1 and Exg1. Several non-canonically secreted proteins lacking secretion signals were also detected, including enolases (Eno1 and Eno2) and heat shock proteins (Hsp7F, Hsp12, and Hsp26). Enolase, a canonically cytoplasmic glycolytic enzyme, has been previously detected in extracellular vesicles and in the cell wall of *S. cerevisiae*, where it is secreted via putative SNARE-dependent unconventional secretion pathway^16,33-35^. Molecular chaperone Hsp12 has been previously reported to be present in the secretome^36^, while we also identified chaperones Hsp7F and Hsp26. Principal component analysis (PCA) (Fig. 1B), clustered heat map analysis (Fig. 1C), and direct quantitative pair-wise comparisons of the secretome in each stress condition to unstressed WT yeast (Fig. S2) showed diverse large and significant differences in protein abundances. All analyses showed that the strongest change in the secretome was in response to heat or ethanol stress.

**Figure 1.**
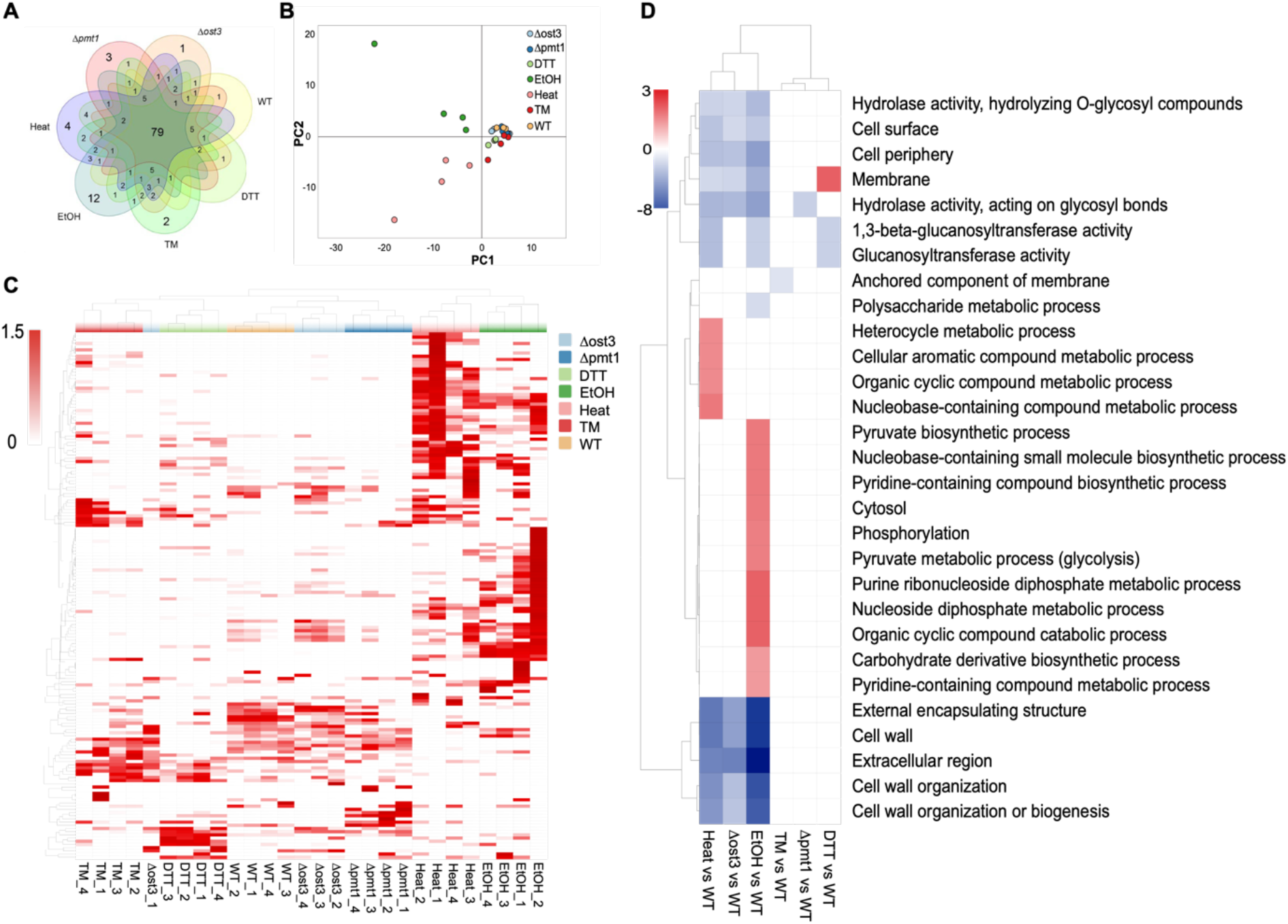
The yeast secretome in response to diverse proteosynthetic stressors. **(A)** Venn diagram of the number of proteins identified in the secretome uniquely or in common under different stress conditions. **(B)** Principal component analysis (PCA) of quantitative secretome profiles under different stress conditions. Each point represents a biological replicate (N=4 for each condition). The variance of PC1=29.7% and PC2=18.1%. **(C)** Clustered heat map of protein abundance in the secretome under different stress conditions (N=4 for each condition). **(D)** Clustered heat map of significantly enriched GO terms identified by GOstats analysis in each comparison. Values shown as -log_2_ of Bonferroni corrected P-value for GO terms which were significantly enriched (*P*<0.05). Values are shown as positive values (red) for GO terms enriched in proteins more abundant in the stress condition and as negative values (blue) for GO terms enriched in proteins less abundant in the stress condition.

To glean insights into the global functional significance of the changes to the secretome in response to stress, we performed GO term enrichment analyses of the proteins with significantly changed abundance in each condition compared to unstressed WT yeast (Fig. 1D and Supplementary Data 5). Ethanol stress showed enrichment of the largest number and most distinct set of GO terms, which were associated with diverse metabolic processes. Strikingly, many proteins associated with glycolysis were significantly more abundant in the secretome of ethanol stressed cells, including Eno1, G3p1, and G3p3 (Fig. S3A-C). Intriguingly, the response in the secretome of Δ*ost3 N*-glycosylation stress and heat stress were similar, with most enriched GO terms for proteins reduced in abundance in the stress conditions being associated with cell wall proteins, such as Gas3 and Gas5 (Fig. S3D-E). Although GO term enrichment was clustered between Δ*ost3 N*-glycosylation stress and heat stress, a clear separation could be observed by PCA (Fig. 1B), suggesting that the changes to the secretome induced by Δ*ost3 N*-glycosylation stress and heat stress compared to unstressed WT yeast were different, but that the affected proteins shared functional roles. Proteins decreased in abundance in the secretome of Δ*ost3*, heat, and ethanol stressed cells all showed GO term enrichment associated with cell wall glycoproteins, including Gas3 and Ccw12 (Fig. S3D and F), consistent with generally decreased canonical secretion under these stress conditions. Reductive DTT stress was associated with increased abundance of membrane-associated proteins in the secretome, such as Tir1 (Fig. S3G), and decreased abundance of proteins with glucanosyltransferase activity, such as Gas3 and Gas5 (Fig. S3D-E). Proteins associated with the GO term “anchored component of membrane” were significantly less abundant in the secretome with tunicamycin stress, including Gas3, Gas5, and Ccw12 (Fig. S3D-F). These proteins also have several *N*-glycosylation sites, consistent with their secretion being impacted by tunicamycin. Δ*pmt1 O*-glycosylation stress triggered a significant decrease in the abundance of proteins associated with hydrolase activity such as Sim1 and Sun4 (Fig. S3H-I), consistent with the long Ser-rich tracts in these proteins that would likely normally be *O*-mannosylated by Pmt1; lack of *O*-mannosylation at these domains may lower the efficiency of protein secretion or the stability of the mature proteins. In addition to proteins associated with enriched GO terms, we also observed changes in some proteins of particular interest. Specifically, the cytoplasmic small heat shock proteins Hsp12 and Hsp26 were more abundant under ethanol and heat stress (Fig. S3K and L). The ER-resident protein disulfide-isomerase Pdi was decreased in abundance under ethanol and heat stress which is consistent with decreased canonical secretion, but increased in abundance under reductive DTT and Δ*ost3 N*-glycosylation stress conditions (Fig. S3L). GO term enrichment analysis revealed a large number of biological activities responding to ethanol stress, especially metabolic processes such as glycolysis, with an increased abundance of glycolytic enzymes in the secretome of ethanol stressed yeast. This was somewhat surprising, given yeast grown in 20% ethanol might be expected to have a reduced dependence on glycolysis.

### Comparison of the stress response in the whole cell and secreted proteomes

The differences in the secreted proteome under the various stress conditions could be due to altered secretion or changes in the whole cell proteome in combination with cell lysis. To discriminate between the possible biological mechanisms underlying the observed strong secretome responses in the various stress conditions, we performed DIA LC-MS/MS proteomic analysis of the whole cell proteome of yeast cells grown in select stress conditions, compared these to the corresponding secreted proteomes, and used GO term enrichment analysis to gain biological insights into the classes of proteins with altered abundance (Suplementary Data 6). For cells grown with ethanol stress, while the secretome showed some proteins with increased relative abundance and others with decreased relative abundance, the vast majority of these were decreased in abundance in the whole cell proteome (Fig. 2A and Supplementary Data 3). Proteins that were reduced in abundance in both the whole cell and secreted proteomes were mostly canonically secreted glycoproteins such as Zps1 (Fig. 2D, G and H), consistent with ethanol stress resulting in lower secretory flux and hence lower glycoprotein abundance in the secretome. The proteins that were increased in relative abundance in the secretome but decreased in abundance in the whole cell proteome under ethanol stress were mainly cytoplasmic glycolytic enzymes such as Eno2 (Fig. 2D, I and J). This is consistent with the key metabolic reprogramming of cells grown in 20% ethanol being a switch from glucose utilisation to ethanol consumption, decreasing the intracellular abundance of these enzymes, but increased cell lysis or UPS in 20% ethanol nonetheless increasing their relative abundance in the secretome.

**Figure 2.**
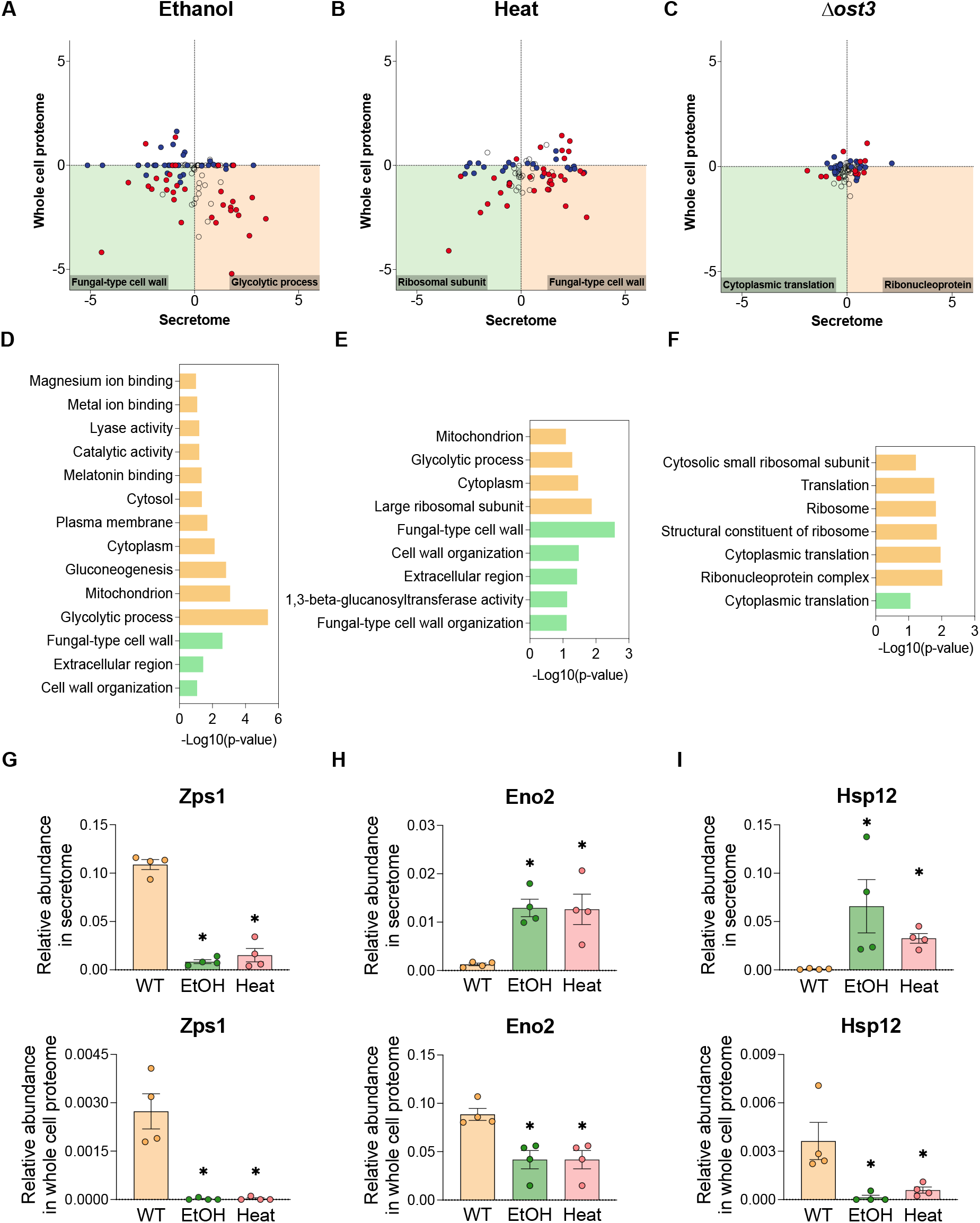
Correlations between stress-induced changes to the secretome and the whole cell proteome. Log_2_(Fold change) in abundance of proteins in the secretome with significant differences in abundance in the stress condition versus Log_2_(Fold change) in abundance of proteins in the whole cell proteome, for **(A)** ethanol stress, **(B)** heat stress, or **(C)** Δ*ost3 N*-glycosylation stress. Red, proteins with significant differences in abundance in the stress condition in both the secretome and the whole cell proteome. Blue, proteins only with significant differences in abundance in the stress condition in the secretome but not in the whole cell proteome. Clear, proteins detected in the secretome but not significantly different in abundance in the stress condition. **(D-F)** Enriched GO terms in each quadrant under ethanol, heat and Δ*ost3 N*-glycosylation stress respectively (Data in Supplementary Data 6). Green, proteins significantly decreased in both the secretome and whole cell proteome, located in the green quadrant of A-C. Orange, proteins significantly increased in the secretome but decreased in whole cell proteome, located in the orange quadrant of A-C. **(G-I)** Individual protein relative abundance in the secretome or whole cell proteome in WT, ethanol, and heat stress. *, significantly different compared to WT (adjusted P<10^-5^).

For cells grown under heat stress, similarly to the response to ethanol stress, the secretome showed proteins with both increased and decreased relative abundance, most of which were decreased in abundance in the whole cell proteome (Fig. 2B). Proteins that were reduced in abundance in both the whole cell and secreted proteomes were mostly cell wall proteins such as Zps1 (Fig. 2E, G and H). The proteins that were increased in relative abundance in the secretome but decreased in abundance in the whole cell proteome under heat stress were mainly heat shock proteins such as Hsp12 (Fig. 2E, K and L). It was particularly surprising that Hsp12 abundance was extremely low intracellularly under ethanol and heat stress, and yet was massively increased in abundance under these conditions in the secretome. Hsp12 is an intrinsically unstructured stress protein that folds upon membrane association and contributes to yeast morphology and membrane stability to protect cells against heat shock^37^. Secretion of heat shock proteins can extend the reach of the stress response into the extracellular microenvironment, where they could stabilize misfolded proteins outside the cell^38^. Δ*ost3 N*-glycosylation stress induced comparatively minor changes to both the secretome and the whole cell proteome (Fig. 2C). Proteins that were reduced in abundance in both the whole cell and secreted proteomes were associated with cytoplasmic translation (Fig. 3F), consistent with the deficiency in *N*-glycosylation induced by Δ*ost3* interfering with productive protein folding and triggering UPR, a key consequence of which is reduced translation ^39^.

**Figure 3.**
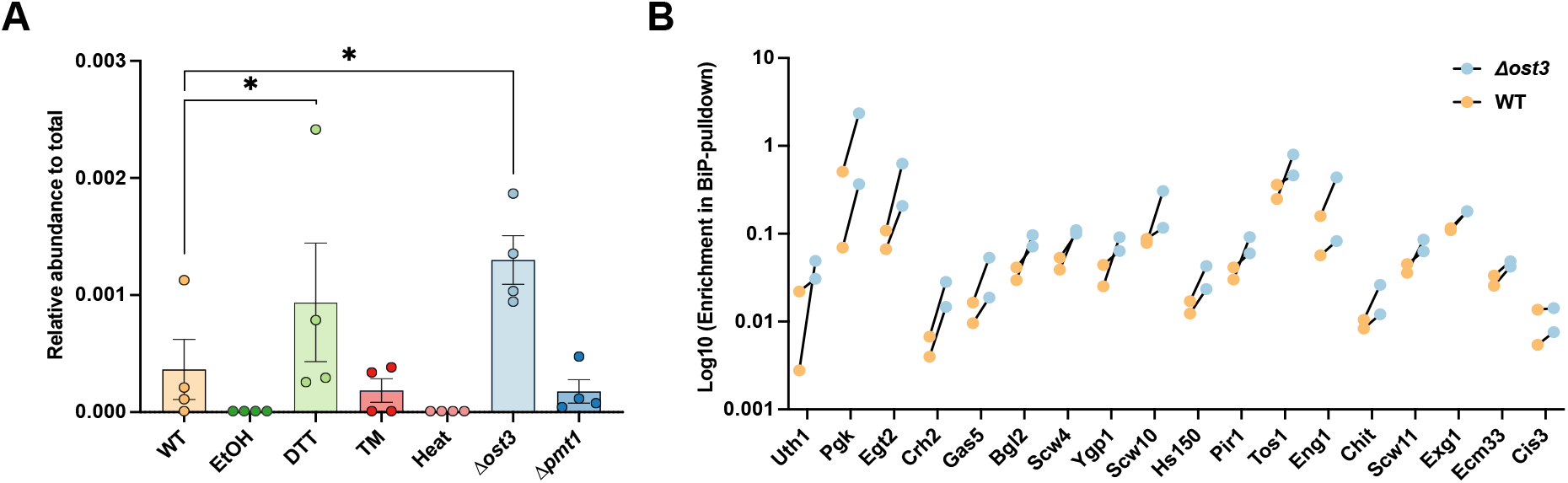
Increased secretion and extracellular physical interaction of ER chaperone BiP with client proteins in ER stress conditions. **(A)** Normalised BiP protein abundance in the secretome under stress conditions. **(B)** Log_10_(Enrichment in BiP-pulldown) from the secretome of Δ*ost3* and WT yeast. N=2, with measurements from each replicate joined by lines.

Cellular stress in Δ*ost3* yeast is driven by inhibition of site-specific *N*-glycosylation and consequent changes in glycoprotein stability and function. It was surprising that heat would cause similar large-scale changes to the secreted proteome as genetic perturbation of *N*-glycosylation, as indicated by similarly clustering based on enriched GO terms (Fig. 1D), and suggested that a key mechanism by which heat affected yeast might also be through inhibition of *N*-glycosylation. To explore this possibility, we mapped site-specific *N*-glycosylation occupancy of cell wall proteins from stressed yeasts under Δ*ost3 N*-glycosylation stress and heat stress, and unstressed WT yeast. We confirmed the large site-specific hypoglycosylation caused by lack of *OST3*^40-42^, but did not detect any measurable change in glycosylation occupancy caused by heat (Fig. S5). It is therefore likely that the similarity in the secreted proteome stress response to heat and *N*-glycosylation deficiency is because both inefficient *N*-glycosylation and high temperatures induce qualitatively similar protein misfolding stress, consistent with the key role of *N*-glycosylation in increasing protein folding efficiency and thermal stability.

In summary, we concluded that in the case of stress caused by heat, ethanol, and Δ*ost3 N*-glycosylation deficiency, the changes to the secretome were caused by a combination of regulated changes to the canonically secreted and whole cellular proteome in response to the particular stress condition, and altered uncontrolled physical lysis or UPS. That is, the yeast secretome reflects the growth conditions, intracellular metabolic and proteomic status of yeast, and quantitatively varies in response to growth under diverse stresses.

### BiP binds secreted proteins under stress

When inspecting the stress-induced secretome, we noticed the presence of canonically intracellular ER chaperone BiP under Δ*ost3 N*-glycosylation stress and DTT reductive stress (Fig. 3A), and cytoplasmic redox chaperones Trx1 and Trx2 under heat stress (Fig. S6). BiP is a key Hsp70 chaperone in the ER that associates with translocating nascent polypeptides and misfolded protein substrates by binding exposed hydrophobic surfaces, and assists productive protein folding in the ER by inhibiting non-productive association of misfolded polypeptides^43^. It can facilitate formation and stabilization of aggregated proteins under stress to maintain the protein folding capability in the ER^44,45^. Thioredoxins act as chaperones to maintain cytoplasmic protein stability in response to oxidative stress conditions^46^. As intracellular chaperones, we were surprised to robustly detect BiP and Trx1/2 in the yeast secretome in specific stress conditions. We found it particularly compelling that the chaperones were detected in the secretome in the stress conditions in which their functions would be most relevant: BiP with DTT reductive stress and Δ*ost3 N-*glycosylation stress, and thioredoxin with heat. As a central chaperone in the ER, BiP maintains protein-folding haemostasis through a catalytic cycle involving ATP binding, ATP hydrolysis, and ADP·Pi release to interact with hydrophobic patches of client proteins while acting as a foldase, holdase, translocase, unfoldase, or disaggregase^47^. We thus speculated that BiP and thioredoxins maintained some activity as chaperones outside of the cell, of particular relevance to the particular stress condition.

To explore the hypothesis that BiP could act as an extracellular chaperone in proteosynthetic stress conditions, we aimed to discover potential client proteins that physically interacted with extracellular BiP. Specifically, we hypothesized that proteins in the Δ*ost3* secretome, but not in the secretome of unstressed WT yeast, would physically interact with BiP. To this end, we created yeast with genomically encoded his-tagged BiP and bound BiP-his protein from unstressed WT yeast to Ni-NTA beads as bait. As prey we used the secretome from either *N*-glycosylation stressed Δ*ost3* or unstressed WT yeast, and used DIA LC-MS/MS proteomics to measure proteins that were specifically enriched in the elution fraction of the Δ*ost3* secretome compared to the elution fraction of the secretome of unstressed WT yeast. We could detect several proteins which specifically interacted with BiP when they were in the Δ*ost3* secretome but not in the unstressed WT secretome, including Tos1, Eng1, Egt2, Exg1, and Pgk1 (Fig. 3B). All these proteins with the exception of Pgk1 are canonical cell wall or secreted glycoproteins. Their interaction with BiP specifically when from the Δ*ost3* secretome is consistent with inefficient *N*-glycosylation in Δ*ost3* cells leading to decreased stability or localised misfolding making them extracellular BiP client proteins. Pgk1 is a cytoplasmic glycolytic enzyme, and was one of the proteins with the largest increase in abundance in the secretome of Δ*ost3* yeast, which likely explains its interactions with BiP even though as a cytoplasmic protein it is never glycosylated. Together, these data confirm that BiP interacts more strongly with diverse proteins in the secretome of Δ*ost3* cells than of unstressed WT yeast, and suggests BiP is a functional chaperone or holdase in the extracellular environment that is particularly important under conditions of ER stress.

### Conclusions

Cellular stresses that interfere with protein secretion can affect yeast physiology, activate the UPR in the ER or chaperones across the cell, and alter canonical and unconventional protein secretion. We systematically profiled the yeast secretome under various stress conditions, which identified distinct proteins and pathways associated with specific stress responses that were a combination of changes to the whole cellular proteome and altered cell lysis. We identified several canonically intracellular molecular chaperones in the secretome under various stress conditions. In particular, we found high levels of BiP in the secretome under Δ*ost3 N*-glycosylation deficiency stress and DTT-induced reductive stress, and showed that BiP physically interacts with potential client proteins extracellularly. This is consistent with secreted BiP functioning as a holdase or chaperone under these stress conditions, aiding in the stabilization and function of client proteins outside the cell.

## Methods

### Yeast strains and growth conditions

*Saccharomyces cerevisiae BY4741* wild type (WT: MATa *his3Δ1 leu2Δ0 met15Δ0 ura3Δ0*), Δ*ost3* (MATa *his3Δ1 leu2Δ0 met15Δ0 ura3Δ0 ost3Δ::kanMX*, Open Biosystems), and Δ*pmt1* (MATa *his3Δ1 leu2Δ0 met15Δ0 ura3Δ0 pmt1Δ::kanMX*, Open Biosystems) were used in this study. Yeast synthetic complete media was used for yeast growth.

Yeast were grown in static culture with a series of stressors as described in Supplementary Table 1, with a starting OD_600nm_ of 0.1. After incubation for four days, the sugar consumption and growth rate of each culture was measured by refractometer (PAL-1, Atago^®^) and cell density meter (Ultrospec 10, Biochrom) at OD_600nm_ respectively. To collect the whole cell and cell well fraction under WT and three cellular stress conditions (Δ*ost3*, heat, and EtOH), yeasts were stationarily grown for two days according to Table S2 (N=4) respectively.

To study the BiP-binding client proteins *in vitro* under targeted stress conditions, a C-terminal his-tag and KanMX cassette (^48^) was appended to the open reading frame of *KAR2*/BiP in *BY4741*. His-BiP expressing yeast was grown at 30 ℃ with shaking at 200 rpm until mid-log phase.

### Sample preparation

Yeast secreted proteins were collected from the supernatant of stationary culture by centrifugation at 4500 rcf for 5 min to remove yeast cells, followed by protein concentration using centrifugal filters (10 kDa, 15 mL, Amicon). Concentrated proteins were precipitated by addition of 1 mL of methanol/acetone (1:1 v/v) and incubation at -20 ℃ overnight, followed by centrifugation at 18000 rcf for 10 min and removal of the supernatant. Yeast whole cell proteins and cell wall proteins were extracted as described^49,50^.

His-tagged BiP was purified from the microsomal fraction of yeast, prepared as previously described^51^. Protein purification and pull-down assays were performed as described^52^. Briefly, 500 μL resolubilised microsomal protein extract was mixed and incubated with 250 μL Ni-NTA resin (HisPur™ Ni-NTA Resin, Thermo scientific) at 4 ℃ for 1 h on a rotator, followed by aspiration on gravity-flow columns and washing five times with 500 μL wash buffer (20 mM sodium phosphate pH 7.4, 300 mM NaCl, 25 mM imidazole, and 0.1% Triton X-100). Concentrated secreted proteins from WT and Δ*ost3* cultures were buffer exchanged into 500 μL equilibration buffer (20 mM sodium phosphate pH 7.4, 300 mM NaCl, 10 mM imidazole, and 0.1% Triton X-100) and used separately as prey proteins. 10 μL of prey proteins were precipitated by 500 μL methanol/acetone (1:1 v/v) and incubated overnight at -20 ℃. The remainder of the prey protein samples were mixed with the Ni-NTA resin bait and incubated at 4 ℃ for 1 h on a rotator, followed by aspiration on gravity-flow columns and washing five times with 500 μL wash buffer. BiP with bound proteins was eluted from the resin by 500 μL elution buffer (20 mM sodium phosphate pH 7.4, 300 mM NaCl, 250 mM imidazole, and 0.1% Triton X-100). Eluted proteins were precipitated by addition of 1 mL methanol/acetone (1:1 v/v) and incubation overnight at -20 ℃, followed by centrifugation at 18000 rcf for 10 min and removal of the supernatant.

Precipitated protein samples were reduced/alkylated, digested with trypsin, and prepared for mass spectrometry proteomic analysis as described^53^. Cell wall extracted proteins were prepared for mass spectrometry glycoproteomic analysis as previously described^49,50^.

### Mass spectrometry

Secretome samples were analysed using a Prominence nanoLC system (Shimadzu) and TripleTof 5600 mass spectrometer with a Nanospray III interface (SCIEX) via data dependent acquisition (DDA) and data independent acquisition (DIA) analysis respectively as described^54^. Approximately 0.5–2 μg peptides were desalted on an Agilent C18 trap (300 Å pore size, 5 μm particle size, 0.3 mm i.d. × 5 mm) at a flow rate of 30 μL/min for 3 min, and then separated on a Vydac EVEREST reversed-phase C18 HPLC column (300 Å pore size, 5 μm particle size, 150 μm i.d. × 150 mm) at a flow rate of 1 μL/min. Peptides were separated with a gradient of 10–60% buffer B over 45 min, with buffer A (1% acetonitrile and 0.1% formic acid) and buffer B (80% acetonitrile with 0.1% formic acid). Whole cell protein, cell wall protein, and BiP-binding protein samples were analysed on Zeno TOF 7600 instrument (SCIEX) via DIA analysis as described^55^. Approximately 500 ng of desalted peptides were separated using reversed-phase chromatography on an M-class UPLC system (Waters). Peptides were separated on a NanoEase HSS T3 column (100 Å pore size, 1.8 μm particle size, 300 μm i.d. × 150 mm) (Waters) at a flow rate of 5 μL/min set at 40 °C, with LC conditions as follows: 0–1 min = 3% buffer B (0.1% formic acid in acetonitrile), 1–46 min = 3–45%, 46–52 min = 45– 97%, held at 97% buffer B for 4 min followed by re-equilibration for 4 min with buffer A (0.1% formic acid in water). Eluted peptides were directly analysed on a ZenoTof 7600 instrument (SCIEX) using an OptiFlow Micro/MicroCal source. For data-independent acquisition (DIA), an MS TOF scan across 400–1500 m/z was performed (0.1 s). For MS2, variable windows spanning 399.5–750.5 *m/z* were chosen for fragmentation (0.013 s), with fragment data acquired across 140–1750 *m/z* with Zeno pulsing on. Dynamic collision energy was used. Sample injection order was randomised. All raw data was acquired using SCIEX OS (v2.1.6).

### Data analysis

Peptides were identified using ProteinPilot™ v5.0.2 (SCIEX) for DDA data with standard settings: Sample type, identification; Cysteine alkylation, iodoacetamide; Instrument, TripleTof 5600; Species, none; ID focus, biological modifications; Enzyme, Trypsin; Search effort, thorough ID. Protein identification was searched against the UniProt reference proteome for *Saccharomyces cerevisiae* UP000002311. False discovery rate analysis using ProteinPilot was performed. Peptides were identified using DIA-NN v1.8^56^ for DIA data against searching parameters: fixed modification, propionamide (71.037114, C); variable modifications, Deamidation (0.984016, N); Missed cleavages, 1; enzyme (cut), trypsin (K*, R*,! *P). Peptides and proteins were quantified using PeakView v2.2 (SCIEX). Peptide and protein abundances were re-calculated with false-discovery rate (FDR) cut-off of 1% ^57^. Protein abundances were normalised to the total protein abundance in each sample. For statistical pair-wise comparisons, the PeakView output was reformatted for use with MSstats ^58^. Differential protein abundance was compared using a mixed linear model using MSstats v2.4 ^59^ in R with a significance threshold of *P*<10^-5^. Gene ontology (GO) term enrichment of secretome peoteomics was performed by GOstats v2.39.1 in R with significance threshold of adjusted *P*<10^-5^. Gene ontology (GO) term enrichment of comparison between secretome and whole cell proteome was performed using the online resource https://david.ncifcrf.gov/ with significance threshold of *P*<0.05^60,61^.

## Supporting information

Supplementary Data 1 - Secretome DDA ProteinPilot searching output

Supplementary Data 2 - Secretome SWATH Peakview quantification output

Supplementary Data 3 - FDR-filter recalculated secretome quantification output

Supplementary Data 4 - Normalised secretome quantification results

Supplementary Data 5 - Secretome MSstats and GOstats

Supplementary Data 6 - Secretome and WCE GO enrichment

## Data availability

Mass spectrometry data have been deposited to the ProteomeXchange Consortium (http://proteomecentral.proteomexchange.org) via the PRIDE partner repository^62^ with the dataset identifier PXD063255.

## Acknowledgements

We thank Dr Amanda Nouwens and Peter Josh at The University of Queensland, School of Chemistry and Molecular Biosciences Mass Spectrometry Facility for their assistance and expertise.

## Author contributions

S.L. and B.L.S. designed the experiments. S.L. performed the experiments. S.L. and B.L.S. analysed the data. S.L. and B.L.S. wrote the manuscript. All authors read and approved the final manuscript.

## Competing interests

The authors declare no competing interests.

## Notes

### Competing Interest Statement

The authors have declared no competing interest.

